# Integrating projected habitat change and existing population trends to prioritise conservation, monitoring, and research for Indian birds

**DOI:** 10.1101/2024.11.27.625604

**Authors:** Vishesh L. Diengdoh, Umesh Srinivasan

## Abstract

Understanding species’ vulnerability to global change requires approaches that integrate projections of future environmental change with empirical population trends; such integrated assessments remain limited in biodiverse, data-limited tropical regions. In this study, we develop and apply a scalable, decision-oriented framework that synthesises climate and land-cover projections with existing contemporary population data to assess species-level vulnerability and conservation, monitoring, and research priorities for Indian birds. Using an ensemble of machine-learning habitat-suitability models, we projected current and future habitat suitability for 955 species under two 2100 scenarios (SSP3–7.0 and SSP5–8.5). We then integrated these projections with population trends from the State of India’s Birds assessment. We found habitat responses were highly heterogeneous: 507 species were projected to lose suitable habitat under both future scenarios, 327 species were projected to gain habitat, and 121 species exhibited mixed or uncertain responses. The direction and magnitude of projected change varied substantially within and among habitat-specialisation guilds, with overall losses exceeding gains. The synthesis of habitat projections with population trends revealed 121 (mostly wetland) species facing ongoing population decline and future habitat loss, representing high-priority targets for immediate conservation action. A further 300 species, largely forest-associated, were projected to lose habitat but lacked reliable population trend data, highlighting substantial monitoring gaps, while species with mixed or uncertain habitat trajectories emerged as key research priorities. By explicitly integrating projected habitat change, contemporary population trajectories, and data availability, this study advances a transferable framework for prioritising conservation action, monitoring, and research under accelerating global change.

## Introduction

Understanding and anticipating species’ responses to global change is a central challenge in conservation biology. Species distribution models (SDMs) are widely used to project how climate and land-cover change may reshape species’ future distributions (Araújo and Peterson 2012; Powers and Jetz 2019), while population estimations provide empirical evidence of ongoing demographic change (Neate-Clegg et al. 2020; SoIB 2023; Viswanathan et al. 2025). However, future distributions and population trends are still rarely combined within a single, species-specific analytical framework that directly informs conservation prioritisation (Pacifici et al. 2015; Visconti et al. 2016; Spooner et al. 2018). Although some studies have coupled habitat projections with demographic or population-viability models to explore extinction risk under future scenarios (e.g., Keith et al. 2008), such approaches remain uncommon at large taxonomic scales and regions. Thus, conservation planning often relies either on forward-looking projections that lack demographic context or on retrospective population trends that do not account for future environmental change, limiting the ability of practitioners to prioritise actions under conditions of rapid and uncertain change (Jetz et al. 2019).

Birds provide an ideal taxonomic group for addressing this gap. They are well distributed across biomes and play important roles in ecosystem functioning (BirdLife International 2022). However, they are threatened by the synergistic effects of several factors including land cover and climate change which drive declines in species richness, abundance, habitat, and distribution (Brook et al. 2008; Oliver and Morecroft 2014; Haddad et al. 2015; Scheffers et al. 2016; Davison et al. 2021). Globally, nearly half of all bird species show population declines (BirdLife International 2022) highlighting the urgency of identifying species and communities most vulnerable to ongoing and future environmental change.

India represents a particularly important but underexamined context for integrative assessments of global change impacts and conservation prioritisation. India is a megadiverse country that houses 1,358 bird species (Maheswaran and Alam 2024) and recent estimates indicate 142 species have declining populations (SoIB 2023; Viswanathan et al. 2025). Overall, national conservation planning remains strongly forest-centric even as growing evidence indicates that open habitats and wetlands are highly threatened (Joshi et al. 2018; Lahiri et al. 2023; Ahmad et al. 2024; Kundu et al. 2024).

While global assessments have examined how climate and land-cover change influence bird distributions (e.g., Foden et al. 2013; Powers and Jetz 2019; Beyer and Manica 2020), such large-scale syntheses can obscure regional variation, masking threats that are acute in biodiverse, rapidly changing tropical nations. Conversely, species-specific case studies (e.g., Tamang et al. 2023; Lele et al. 2024), though detailed, limit broader comparative inference due to restricted taxonomic scope and inconsistent methods. Consequently, there remains a critical need for a standardised, national-scale assessment that evaluates how global change will affect entire avian communities within a single country.

To address these needs, we assess the impacts of land-cover and climate change on habitat suitability for Indian bird species under present conditions (2025) and two future scenarios (2100: SSP3–7.0 and SSP5–8.5). Specifically, we (i) quantify projected changes in suitable habitat for 955 species, (ii) examine how the magnitude and direction of change vary across habitat-specialisation guilds (alpine and cold desert, forest, forest and plantation, grassland, grassland and scrub, non-specialised, open habitat, and wetland), and (iii) integrate projected habitat trajectories with contemporary population data from the State of India’s Birds (SoIB 2023; Viswanathan et al. 2025). This integration allows identification of species experiencing both current population decline and future habitat loss, species projected to lose habitat but lacking population trend data, and species with mixed or uncertain responses. By combining future habitat projections with national-scale demographic information, we provide a generalisable framework for identifying conservation, monitoring, and research priorities across India’s avifauna and for supporting evidence-based biodiversity planning in rapidly changing tropical regions.

## Methods

### Species

Species presence records were downloaded from Global Biodiversity Information Facility (GBIF, 15 October 2023). Records were restricted to ‘human observation’ entries from the ‘eBird Observation’ dataset within India and to observations collected from 1981 onwards. Duplicate records were removed and spatial autocorrelation was reduced by retaining a single record per 1-km pixel. To minimise errors, records occurring in land-cover classes considered inconsistent with each species’ documented habitat specialisation were excluded.

Pseudo-absences were generated using a target-species method (Phillips et al. 2009) i.e., the presence of other species was used as pseudo-absences. These were at least 1000 m from each other and presence points. The number of pseudo-absences generated was equal to the number of presences.

After data cleaning and thinning, only species with at least 100 data points were considered for analysis because it provided data adequate for robust model training and testing.

For each species, the study area was limited to its IUCN range within India.

### Predictors

Habitat suitability was modelled using CHELSA bioclimatic variables (Karger et al. 2017, 2018), land cover (water, forest, grassland, barren, cropland, urban, and permanent snow and ice; Chen et al. 2022) and Shuttle Radar Topography Mission (SRTM) elevation (Reuter et al. 2007; Jarvis et al. 2008) as predictors.

The bioclimatic variables for the periods 1981–2010 and 2011–2040 (SSP3–7.0, selected as the scenario most consistent with recent observed trends) were used to represent present climatic conditions; species presence and pseudo-absence records temporally matched these periods. Future climate projections were derived from the period 2070–2100 under SSP3–7.0 and SSP5–8.5 across five general circulation models (GCMs). Land-cover data for 2020 (SSP3–7.0) represented present conditions, while projections for 2100 under SSP3–7.0 and SSP5–8.5 represented future land-cover scenarios.

For each species, multicollinearity among numeric predictors was reduced by removing highly correlated variables (≥ 0.7) using a stepwise variance inflation factor (VIF) function in the *usdm* R package (Naimi et al. 2014).

### Habitat suitability model

To build the habitat suitability models, the data was split into 70% training and 30% testing sets.

Model optimisation and threshold selection were conducted using cross-validation on the training data via the *caret* R package (Kuhn 2008). Four machine-learning algorithms representing different modelling approaches were implemented: artificial neural networks, gradient boosting machines, k-nearest neighbour, and random forest. The trained and optimised algorithms were used for projecting habitat suitability under present conditions and future scenarios; these outputs were averaged (unweighted) to result in the ensemble habitat suitability model.

The accuracy of the cross-validated models were assessed using the Area Under the Curve (AUC) and True Skill Statistic (TSS). Similarly, AUC and TSS were calculated for the (present condition) ensemble models using the testing data.

The thresholds for converting probabilistic (0.0–1.0) ensemble projections to binary (0/1) were obtained using the sensitivity–specificity method (Liu et al. 2005), averaged across cross-validation runs.

All analyses were conducted in R v4.4.0 (R Core Team 2025).

## Results

Habitat suitability models were developed for 955 bird species under present conditions and two future scenarios. Model performance was high, with minimal differences between cross-validated and ensemble Area Under the Curve (AUC) and True Skill Statistic (TSS) values (Supplementary Information 1).

Across all species, 507 (53.08%) were projected to lose suitable habitat under both future scenarios, 327 (34.24%) were projected to gain suitable habitat, and 121 (12.67%) exhibited mixed or uncertain responses (Table 1; SI 1). These patterns varied substantially across habitat-specialisation guilds, with both projected “winners” and “losers” present in all guilds. Most guilds exhibited a higher proportion of species projected to lose habitat; the exceptions were forest and open-habitat guilds, which contained proportionally more species projected to gain habitat.

**Table 1.** The number (percentage) of species projected to experience decreases, increases, or mixed changes in suitable habitat across habitat-specialisation guilds.

| Habitat specialisation | Number (percentage) of species |  |  |
| --- | --- | --- | --- |
|  | Decrease | Increase | Mixed |
| Alpine & Cold Desert | 21 (72.4) | 1 (3.4) | 7 (24.1) |
| Forest & Plantation | 93 (64.6) | 33 (22.9) | 18 (12.5) |
| Forest | 113 (35.5) | 162 (50.9) | 43 (13.5) |
| Grassland & Scrub | 11 (55.0) | 4 (20.0) | 5 (25.0) |
| Grassland | 13 (65.0) | 5 (25.0) | 2 (10.0) |
| Non-specialised | 108 (65.1) | 37 (22.3) | 21 (12.7) |
| Open Habitat | 35 (42.2) | 42 (50.6) | 6 (7.2) |
| Wetland | 113 (64.6) | 43 (24.6) | 19 (10.9) |

The magnitude and direction of projected changes in suitable habitat differed strongly among habitat guilds (Fig. 1). Among species projected to lose habitat (Fig. 1a), the largest median declines were observed in grassland-and-scrub, non-specialised, and wetland guilds, with reductions ranging from 10.8 to 14.1 million hectares between present conditions and SSP5–8.5. In contrast, forest species exhibited the smallest median declines, averaging approximately 0.93 million hectares. Among species projected to gain habitat (Fig. 1b), the largest median increases occurred in open-habitat and wetland guilds, with expansions of approximately 6–12 million hectares by SSP5–8.5. The smallest gains (<1 million hectares) were projected in alpine–cold desert, forest, and non-specialised guilds.

**Fig. 1.**
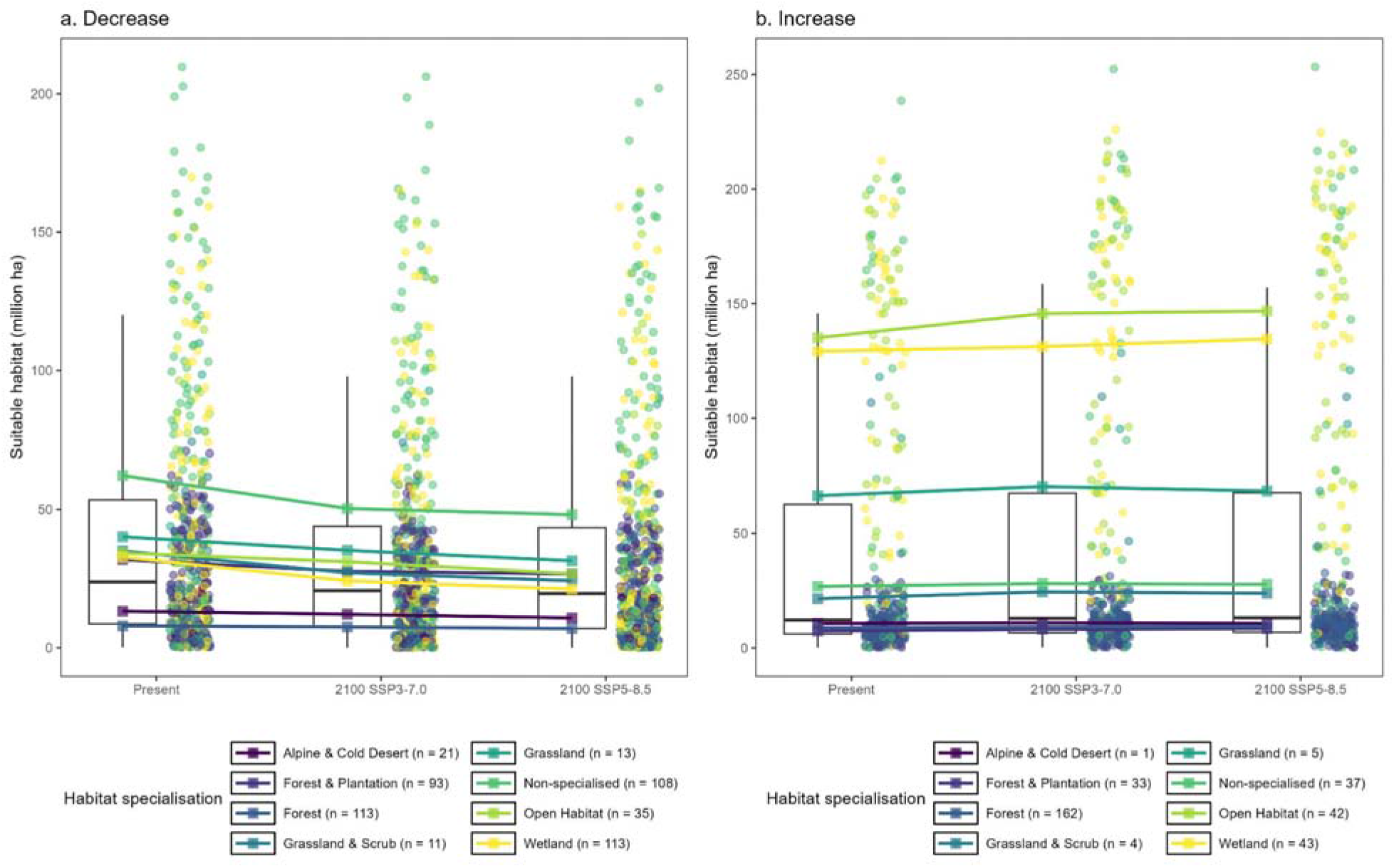
Boxplots showing suitable habitat area (million ha) under present and future scenarios for species projected to (a) decrease and (b) increase in suitable habitat. The square points indicate guild-level medians and round points indicate individual species. Species with mixed responses are not shown.

Integrating projected habitat trajectories with population trends from the State of India’s Birds assessment (SoIB 2023; Viswanathan et al. 2025) identified three groups of species of conservation concern (Fig. 2; SI 1). A total of 121 species, predominantly wetland-associated, were characterised by both ongoing population declines and projected future habitat loss, were identified as the conservation priority group. The monitoring priority consisted of 300 species, largely forest-associated, were projected to lose habitat but lacked reliable population trend data. Finally, 114 species, again primarily forest-associated, exhibited mixed or uncertain habitat projections in combination with declining or data-deficient population trends, representing a research priority.

**Fig. 2.**
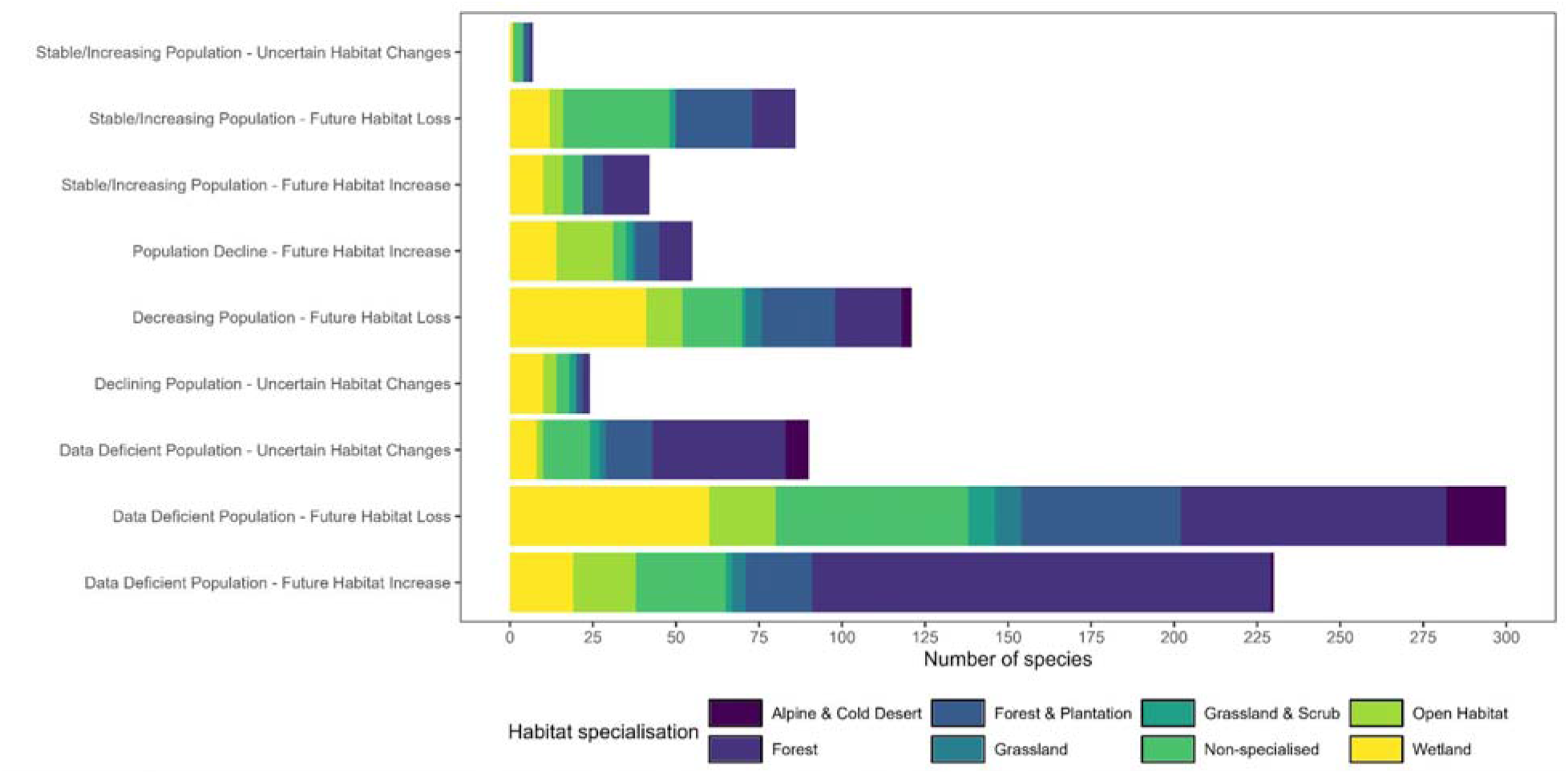
Stacked bar chart showing number of species assigned to conservation priority (declining populations with future habitat loss), monitoring priority (data-deficient populations with future habitat loss), and research priority (mixed habitat responses with declining or data-deficient population trends).

## Discussion

Our study provides one of the most comprehensive national-scale assessments to date of how projected changes in climate and land cover could alter habitat suitability for Indian bird species and how these projections intersect with contemporary population trajectories. Habitat-suitability models revealed heterogeneous responses to global change, with a greater proportion of species projected to lose suitable habitat than gain it. These responses varied substantially both between and within habitat-specialisation guilds, with grassland–scrub, non-specialised, and wetland species experiencing the largest median declines, and forest-associated species showing comparatively smaller declines. Integrating projected habitat trajectories with empirical population trends allowed identification of three distinct priority groups: species facing concurrent population declines and future habitat loss, species projected to lose habitat but lacking reliable trend data, and species with mixed or uncertain habitat responses combined with ambiguous trend information.

Our findings are broadly consistent with global evidence demonstrating heterogeneous species vulnerability to climate and land-cover change, mediated by species-specific traits, habitat associations, and regional context (Foden et al. 2013; Pacifici et al. 2015; Urban et al. 2016; Dyderski et al. 2018; Powers and Jetz 2019; Tuan et al. 2023). The comparatively small median loss of suitable habitat projected for forest-associated species could be due to the buffering effects of forest microclimates (Bonan 2008; De Frenne et al. 2021). In the land-cover scenarios applied here, projected change between 2020 and 2100 was substantially lower for forest classes (13%) than for grassland classes (48%), which may partially explain the more muted responses of forest species. In contrast, larger projected losses for open-habitat and wetland species are consistent with long-documented patterns of habitat conversion and degradation in these systems (Bardgett et al. 2021; Fluet-Chouinard et al. 2023; Lahiri et al. 2023; Lyons et al. 2023) and with empirical evidence linking grassland habitat loss to population declines of grassland birds in India (Ramesh et al. 2025).

By synthesising projections of future climate and land-cover change with contemporary population trends for Indian birds, our study provides the type of forward-looking, evidence-based assessment explicitly called for in India’s National Wildlife Action Plan 2017–2031 (NWAP; MoEFCC 2017) and the Kunming–Montreal Global Biodiversity Framework (GBF; Stephens 2023) both of which emphasise integrating climate change and accessible biodiversity knowledge into conservation planning. The identification of conservation, monitoring, and research priority species directly supports national and global commitments to halt species declines, including Target 4 of India’s updated National Biodiversity Strategy and Action Plan 2024-2023 (NBSAP; MoEFCC 2024) and GBF Target 4, as well as NWAP objectives on strengthening research and long-term monitoring. Together, these outputs provide a coherent, policy-aligned evidence base that strengthens India’s national biodiversity strategy, supports international reporting obligations, and offers a generalisable framework for conservation planning under accelerating global change.

Our study has limitations that should be considered when interpreting the results. First, the 1-km spatial resolution of environmental predictors may obscure fine-scale heterogeneity and microrefugia. Second, biases in citizen-science occurrence data (GBIF/eBird) may influence projections, despite efforts to mitigate bias. Third, our correlative models do not explicitly incorporate biological processes such as dispersal, adaptation, species interactions, and other stressors (e.g., pollution) that may alter realised responses (Araújo and Peterson 2012; Dormann et al. 2012). Future research should integrate mechanistic and demographic models, use fine-grain environmental data where available, and develop species-specific life-history models that can capture adaptive capacity and local population dynamics (Pacifici et al. 2015; Urban et al. 2016).

In conclusion, this study delivers a scalable, transparent framework for integrating projected habitat suitability with contemporary population dynamics to prioritise conservation action, monitoring, and research for Indian birds. By moving from projection to prioritisation, the framework enables evidence-based decisions in the face of accelerating climate and land-use pressures. As global change continues to intensify, such integrative approaches will be essential for directing limited conservation resources toward the species, habitats, and data gaps where they can have the greatest impact.

